# sciMET-cap: High-throughput single-cell methylation analysis with a reduced sequencing burden

**DOI:** 10.1101/2023.07.12.548718

**Authors:** Sonia N. Acharya, Ruth V. Nichols, Lauren E. Rylaarsdam, Brendan L. O’Connell, Theodore P. Braun, Andrew C. Adey

**Author notes:** These authors contributed equally to this work (Sonia N. Acharya, Ruth V. Nichols).

## Abstract

DNA methylation is a key component of the mammalian epigenome, playing a regulatory role in development, disease, and other processes. Robust, high-throughput single-cell DNA methylation assays are now possible (sciMET); however, the genome-wide nature of DNA methylation results in a high sequencing burden per cell. Here, we leverage target enrichment with sciMET to capture sufficient information per cell for cell type assignment using substantially fewer sequence reads (sciMET-cap). Sufficient off-target coverage further enables the production of near-complete methylomes for individual cell types. We characterize sciMET-cap on human PBMCs and brain (middle frontal gyrus).

## Background

DNA methylation is a key component of the mammalian epigenome with crucial roles in gene regulation, lineage specification, and genome organization. Aberrant methylation patterns are characteristic of a multitude of disease states including cancer [1–6]. Like transcription and other epigenetic properties, cell type heterogeneity necessitates analysis of DNA methylation at the single-cell level to achieve cell type or cell state deconvolution in complex tissues, but the ability to assess methylation has lagged behind single-cell RNA transcription or chromatin accessibility assays with respect to cell throughput capabilities. This has largely been due to two factors: the lack of a robust technology that is capable of producing large numbers of single-cell methylomes; and the cost of sequencing genome-wide methylation profiles. The cell throughput challenge is due to the way in which DNA methylation is assessed, using either chemical (bisulfite) or enzymatic methods to convert unmethylated cytosine bases to uracil, which are in turn sequenced as a thymine while methylated cytosine bases retaining their identity. Both techniques for this conversion require DNA to be denatured and involve multiple washes and buffer exchanges incompatible with many standard high-throughput approaches, such as droplet-based techniques. This has forced single-cell methylation technologies to stick to a plate-based format where each individual cell undergoes the conversion process within a single well of a plate [7–11]. While this approach can be scaled to great cell numbers using robotics and large quantities of microwell plates [12–14], it is not a viable strategy for the large majority of research programs.

We recently described an improved version of a high-throughput technique for single-cell methylation analysis [15], sciMETv2, which provides improved usability, cell coverage, and robustness when compared to the original sciMET workflow [16]. This technology leverages combinatorial cell barcoding where nuclei are fixed and carried through nucleosome disruption prior to indexed tagmentation using fully-methylated adapters. The nucleosome disruption step enables genome-wide access for the tagmentation reaction providing global coverage [17,18]. Nuclei are then pooled and distributed to new wells with a limiting dilution or sorting such that the probability of two nuclei with the same tagmentation index ending up in the same well is low. Bisulfite-based conversion is then performed on the pool of cells in each well, amortizing both the labor and costs over multiple cells and enabling the production of datasets comprised of thousands of cells in a single experiment.

The sciMETv2 workflow addresses the first challenge regarding throughput for single-cell methylation analysis; however, the costs associated with sequencing are not reduced, with a typical read count ranging from two to six million raw reads per cell [14,15,19]. Such high read counts are necessary as at least 300-500 thousand CG dinucleotides must be captured genome-wide per cell in order to perform cell type assignment [15]. The issue of high sequencing requirements is also a factor in bulk cell assays, which prompted the development of targeted techniques such as reduced representation bisulfite sequencing (RRBS), which leverages restriction digests to target a subset of the genome. However, this approach misses a substantial portion of putative regulatory regions [20,21]. Alternatively, hybridization capture using probes to known regulatory loci has emerged as a means of enriching for all regulatory regions of interest, with platforms that are compatible with post-conversion libraries as opposed to enrichment prior to bisulfite conversion [22]. Furthermore, the enrichment techniques allow for multiple barcoded libraries to be pooled prior to capture, suggesting that it may be compatible with the barcoded single-cell libraries produced by sciMET.

Here, we leverage a post-bisulfite methylome capture panel targeting 123 Mbp of regulatory regions that is commercially-available from Twist Bioscience to enrich libraries generated using sciMETv2.LA and sciMETv2.SL techniques (Fig. 1a-b). We systematically assess the performance of cluster resolution and differentially methylated region (DMR) detection using a ∼2,000 cell sciMETv2.SL preparation on human peripheral blood mononuclear cells (PBMCs) for four low-read count datasets (median ∼150,000 raw read pairs) covering three capture conditions and a no-capture control, all using the same input library. We observed universally improved results for all metrics when capture is implemented. We also demonstrate that the accumulation of off-target reads in aggregated clusters enables near-complete methylomes for assessment of loci beyond the target panel. We next demonstrate the production of an additional ∼4,000 cell profiles sequenced on a single run of a benchtop sequencer to produce a combined ∼6,000 cell PBMC sciMET-cap dataset. Finally, we demonstrate capture performance on a sciMETv2.LA preparation on human middle frontal gyrus covering ∼1,000 cells that was deeply sequenced (>7 billion raw reads across the dataset) and previously described [15]. We leveraged sciMET-cap and produced a low-read count dataset (∼340 million raw reads total) which produced comparable clustering results compared to the high-depth non-capture dataset. Altogether, our results suggest that targeted enrichment sequencing of regulatory regions can dramatically reduce sequencing costs while simultaneously improving data quality.

**Figure 1.**
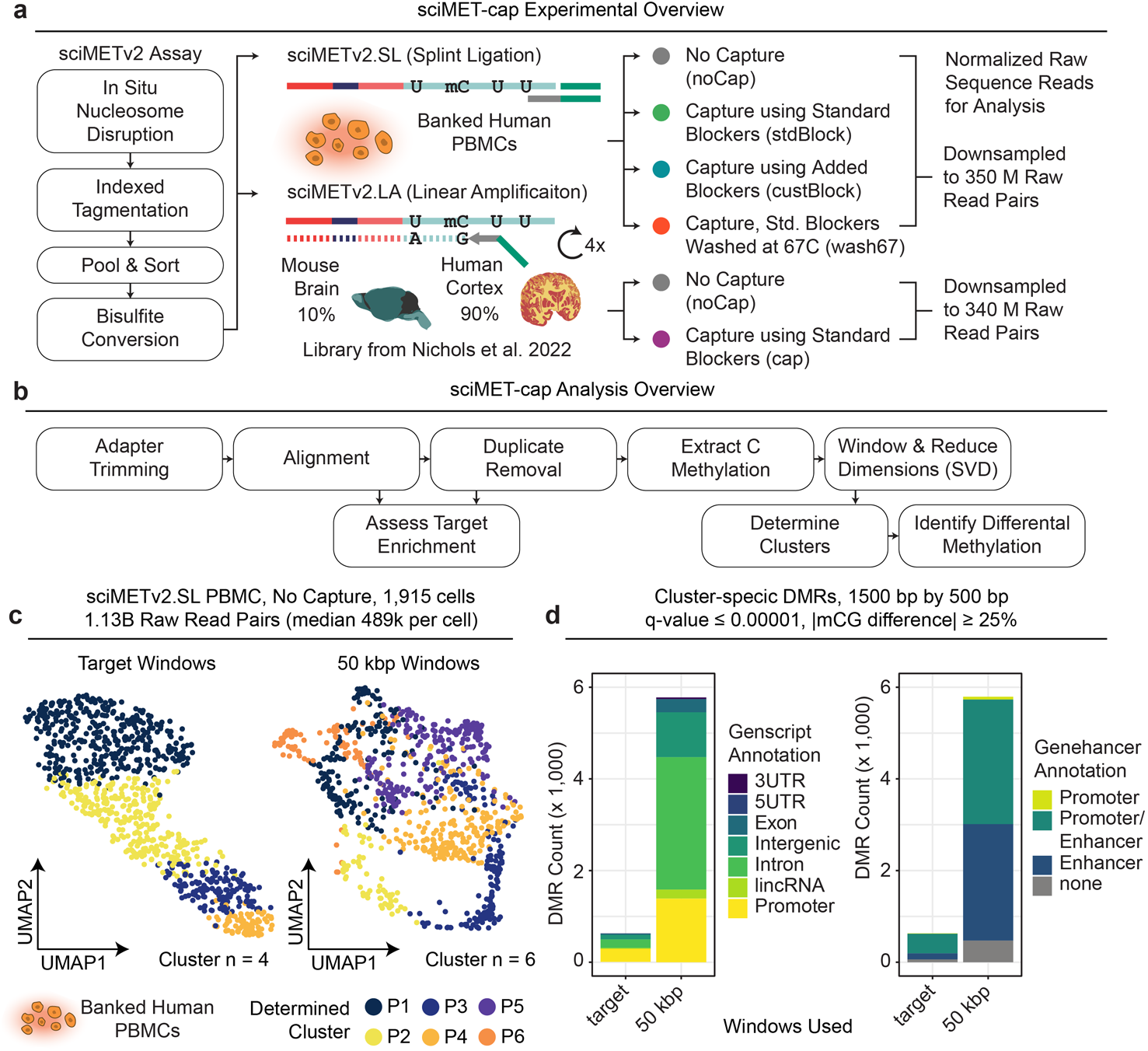
sciMET-cap experimental overview and input PBMC dataset. **a.** Overview of sciMETv2 workflow with the splint ligation (sciMETv2.SL) method used to generate the input PBMC library which was used as input to three capture conditions as well as a no capture control. Sequence data was downsampled to 350M raw read pairs for subsequent analysis. The linear amplification method (sciMETv2.LA) was used to produce a library from mouse brain and human middle frontal gyrus; human cell IDs were used for subsequent comparison between capture methods and a no capture control, both downsampled to 340M raw read pairs. **b.** Overview of analyses used to assess dataset performance. **c.** Clustering and UMAP visualization of cells from the input PBMC dataset using two different window sets. **d.** Cluster-specific differentially methylated regions (DMRs) called for each windowing method broken down by Genscript and Genehancer annotations.

## Results

### sciMETv2 performance on peripheral blood mononuclear cells

To develop sciMET-cap we produced a sciMETv2 library using the rapid, splint ligation (sciMETv2.SL) workflow on a human peripheral blood mononuclear cell specimen (PBMC). This preparation performed comparably to previously-published sciMETv2.SL data generated on human cortex, achieving a median of 283,317 unique CG sites assessed for 1,915 cells at a sequencing depth of 489,303 median raw sequence reads per cell (1.13 billion total raw reads). At this depth of sequencing, a median of 90.09% of reads were unique per cell, suggesting additional depth would yield higher CG coverage before reaching diminishing returns. However, this read depth was sufficient to perform cell type clustering using two strategies (Fig. 1c).

We first leveraged the genome-wide nature of the data, using 50 kbp non-overlapping tiles across the main-series chromosomes as previously described [15]. This produced 6 clusters (Fig. 1c), resulting in 5,914 cluster-specific DMRs (Fig. 1d; q-value ≤ 1e-5, |methylation difference| ≥ 25%). We next utilized the set of windows targeted by the Twist Methylome Capture Panel by merging all probe locations across main-series chromosomes (hg38: chr1-22, X, Y) for a total of 636,303 windows spanning 123,043,166 bp (target). We reasoned that these windows, which were designed based on a wealth of bulk-cell methylome literature, represent the most variable regions of the methylome composed of CG islands and shores, tissue-specific differentially methylated regions (DMRs), enhancers, and other putative regulatory elements [22]. This produced a total of 4 clusters without clear separation (Fig. 1c) and a total of 813 cluster-specific DMRs (Fig. 1d).

Importantly, for purposes of comparison between conditions throughout this work, we leveraged the same clustering parameters for consistency. We then aggregated cell profiles within each cluster to perform cluster-specific DMR identification, again using consistent calling parameters and using the same set of 1,500 bp windows that slide by 500 bp. For all DMR analyses, we include every 1,500 bp window that meets significance criteria for each cluster comparison, meaning some DMR windows may overlap and would typically be collapsed in subsequent analyses. However, for comparison purposes, retention of all significant windows provides a more accurate comparison of DMR identification power. Similarly, windows that are called as significant in more than one cluster when compared to all other clusters are also retained for the same reason.

### Evaluation of sciMET-cap target enrichment conditions

The nature of sciMETv2 enables a high final library yield due to the 96-well plate final PCR amplification, providing ample material for sequencing as well as multiple target capture reactions. We therefore leveraged the input PBMC sciMETv2.SL library to perform three target capture tests: capture using the manufacturer’s recommendations with standard blocking oligos and a wash temperature of 63°C (stdBlock), adding in a custom blocking oligo targeting the Tn5 recognition sequence that is present in sciMETv2 libraries and not in typical bisulfite sequencing assays, also with a 63°C wash temperature (custBlock), and finally using standard blockers and an elevated wash temperature of 67°C to increase stringency (wash67). Each capture library was sequenced separately and downsampled to an identical raw sequence read count of 350 million read pairs along with the no capture control (noCap) to enable a controlled assessment of each method’s performance (Fig. S1).

As anticipated, the raw read count per cell across each condition was comparable with a substantial decrease in unique aligned reads for the capture libraries versus non capture (median = 213,723 noCap, 175,255 stdBlock, 175,004 custBlock, 154,192 wash67) due to an overall reduction in library complexity when enriching for on-target fragments. However, the number of unique CG sites covered was improved for capture libraries with a median ranging from 134,432-139,006 versus 93,676 for the non-captured library, resulting in an improved CG capture rate per raw sequence read. Furthermore, the CG sites in capture libraries are enriched for the regions in the panel and therefore may be more informative for cell state due to the variable nature of these loci. We next compared the three capture conditions for overall target enrichment. Across all metrics little difference was observed between the standard blocker and standard + custom blocker conditions, both with a wash at 63°C, suggesting that the addition of blockers specific to the unique sequences present in the sciMETv2 workflow is not necessary. However, the condition with a wash at 67°C exhibited a greater target enrichment of aligned reads with a median of 10.1-fold versus 9.03 and 9.4 for stdBlock and custBlock, respectively. This improvement was eliminated when assessing only unique reads, with enrichment values dropping to a median of 7.4, 7.7 and 7.1-fold for stdBlock, custBlock and wash67, respectively; owing to the decreased library complexity incurred from target enrichment that was more pronounced in the wash67 condition. Taken together, the higher wash temperature produces a modest improvement in reads on target, yet at a substantial cost in the number of unique reads that can be obtained, resulting in less overall information content obtained per cell at the same raw sequencing depth.

### Clustering and DMR identification using sciMET-cap data

For each sciMET-cap condition we next assessed the ability to identify distinct clusters and cluster-specific DMRs using the equivalent raw read count datasets. As with previous assessments we leveraged either the capture target windows or tiling 50 kbp windows and performed dimensionality reduction, clustering and UMAP visualization (Fig. 2a). As anticipated based on the full non-capture dataset, the target windows for the downsampled no-capture control performed poorly, and produced only 4 clusters, while an improved 6 clusters were resolved using 50 kbp tiling windows. Conversely, each of the capture conditions produced greater cluster counts using the target windows. The custBlock condition identified 7 clusters and the stdBlock and wash67 conditions identified 8 clusters using a target windows. However, with 50 kbp tiling windows, all conditions produced 5 clusters.

**Figure 2.**
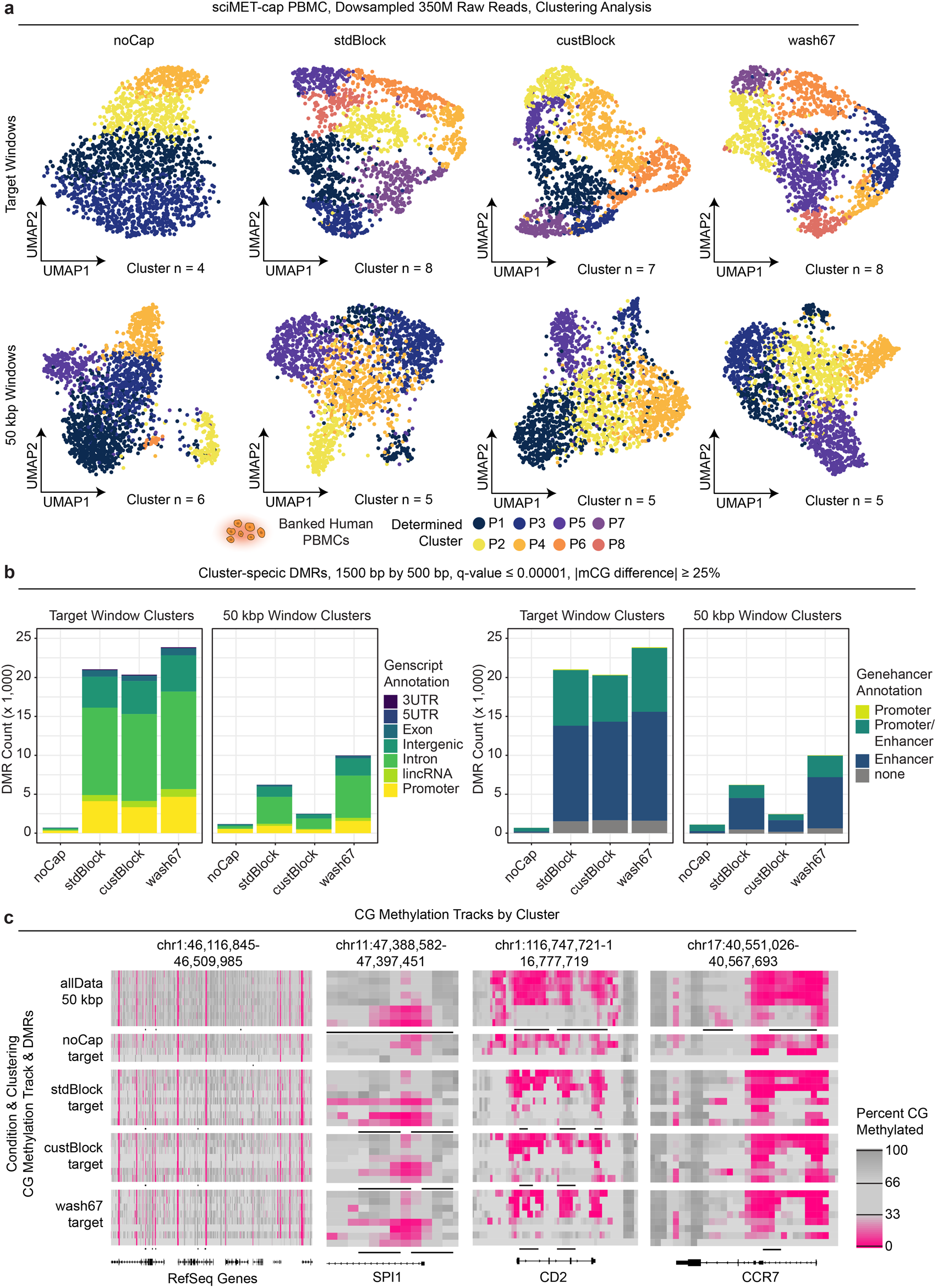
Comparison on sciMET-cap conditions on clustering and DMR analysis. **a.** Clustering and UMAP visualization of each of the four capture conditions and no capture control using either target windows (top) or 50 kbp tiling windows (bottom). **b.** Comparison of cluster-specific DMRs identified across conditions for each window set and colored by Genscript and Genehancer annotations. **c.** Genome browser views of CG methylation levels over 1,500 bp windows tiling by 500 bp split by clusters across conditions for four regions.

To further evaluate clustering performance, we aggregated all read data from each capture experiment as well as the full no capture input sequencing data (Fig. S2a), to produce an aggregate dataset (allData; 1,629,970 median raw reads, 1,169,756 median unique reads, and 551,685 median unique CG sites covered per cell). We then performed clustering with both window strategies, producing 7 clusters using target windows which resulted in 44,034 cluster-specific DMRs; and 8 clusters using 50 kbp windows which resulted in 68,547 DMRs (Fig. S2b). We then compared cluster assignments between each downsampled condition and the allData dataset which produced greater concordance for each of the three capture conditions when compared to the no capture control (Fig. S2c).

We next performed cluster-specific DMR identification for each condition using the downsampled datasets and leveraging both target and 50 kbp window clustering results. Across both clustering sets, the capture conditions produced greater numbers of identified DMRs when compared to the no capture control, at 2,497 (custBlock), 6,224 (stdBlock), and 9,984 (wash67) versus 1,164 (noCap) for the 50 kbp window clustering with a far greater difference for the target window clustering at 20,346 (custBlock), 21,034 (stdBlock), and 23,832 (wash67) versus 694 (noCap). We then compared these DMR calls with those present in the aggregated allData dataset (Fig. S2d). Of the capture experiments, the stdBlock condition identified the largest proportion of DMRs that were present in the allData call sets, with the fewest condition-specific calls. The remaining conditions also identified a majority of DMR calls present in the allData call sets. The no-capture downsampled control (least DMRs) identified a minority of regions from the allData call sets, with some entirely unique regions identified. When evaluating marker gene methylation across clusters, each of the capture datasets produced distinct patterns at cell type specific marker genes in a cluster specific manner. DMRs identified by our prior analysis were frequently found at the promoters of these marker genes, suggesting that our differential methylation analysis is identifying true cell-type-specific patterns of methylation. In contrast, the no capture control lacked specificity, showing a broad pattern of CG methylation across most clusters (Fig. 2c).

### sciMET-cap can produce rich single-cell methylome datasets using a benchtop sequencer

We next sought to demonstrate the throughput of sciMET-cap that can be produced from a single sequencing run on a benchtop sequencer, specifically the Illumina NextSeq 2000 benchtop sequencer capable of producing up to 1.2 billion 100 bp paired reads in a single run. We prepared another sciMETv2 library using the rapid splint ligation workflow on PBMCs from the same individual, targeting approximately 4,000 cells. We then performed a single capture using the standard blockers and blocking conditions and sequenced the library on a single flowcell. This resulted in 4,037 single cell methylomes with a median of 233,400 raw reads per cell, producing a median of 166,270 unique CG sites covered per cell with a unique read target enrichment of 7.2. Overall capture performance was comparable to the prior work for half the cell count, suggesting capture probe saturation was not a factor.

The additional cells were then merged with the non-downsampled stdCap dataset detailed previously, based on the matching preparation and capture conditions to produce a dataset of 5,952 PBMC sciMET-cap profiles. The dataset was then taken through target window clustering and UMAP visualization producing 14 distinct clusters divided into four major groups representing monocytes, B cells, NK cells, and T cells, with two branching groups of clusters representing CD4 and CD8 subsets (Fig. 3a). Cell type assignment was determined by assessing the methylation status of promoter regions at known marker genes and confirmed by comparing profiles to bulk cell WGBS profiles produced on sorted PBMC cell populations as a part of the BLUEPRINT Epigenome project [23] (Fig. 3b). DMR calling produced a total of 195,483 called regions (Fig. 3c) which were achieved using a total raw read count of 1.7 billion – a 1.5-fold increase in raw sequencing when compared to the input PBMC library sequenced without capture that was able to produce only 5,914 DMRs using the same significance criteria, for a 33-fold improvement.

**Figure 3.**
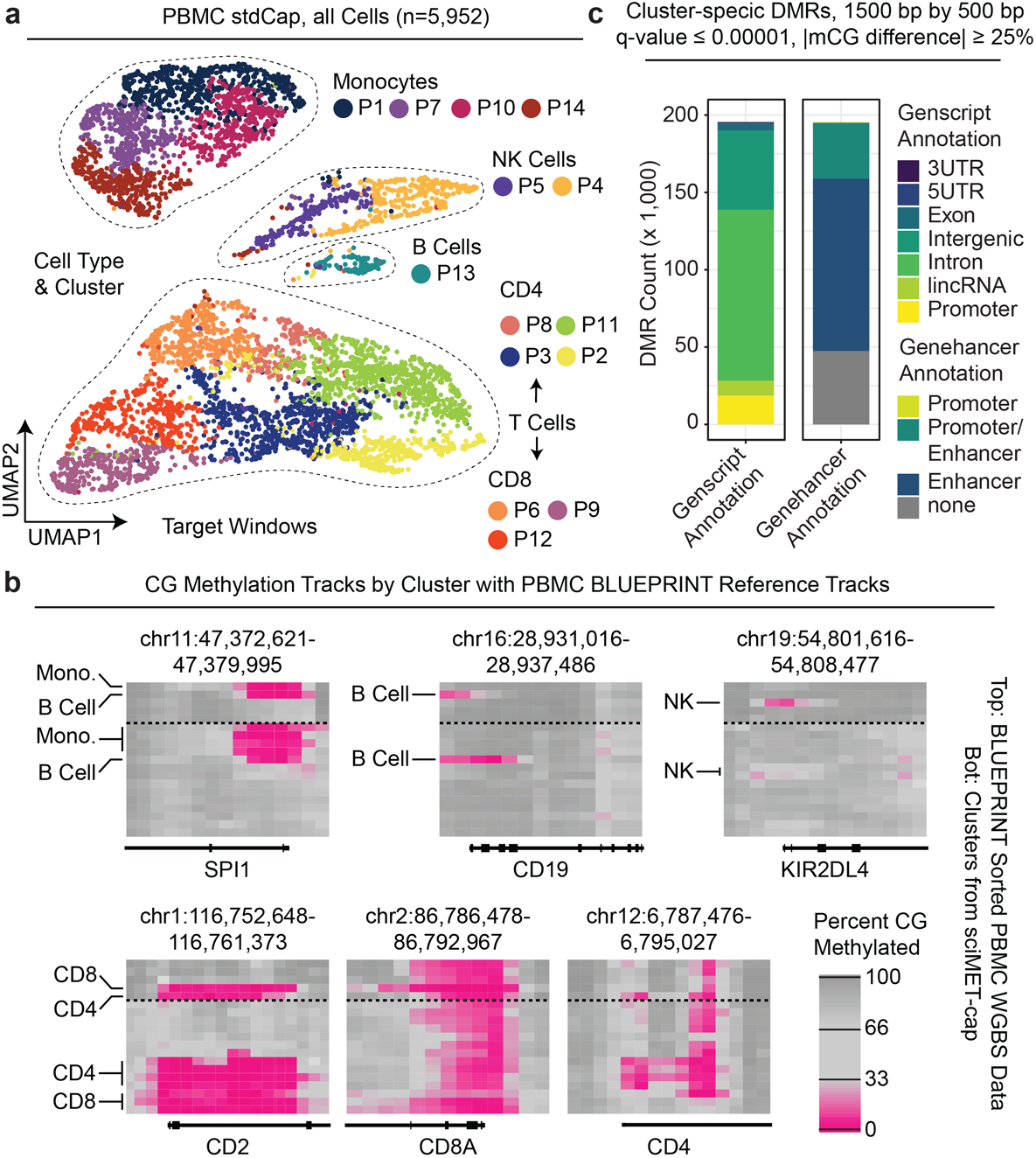
Demonstration of sciMET- cap performance using a benchtop sequencer. **a.** UMAP visualization and cell type clustering of a nearly 6,000 cell sciMET-cap PBMC dataset. **b.** Genome browser tracks of DNA methylation at marker genes. Top: Epigenome Blueprint bulk bisulfite sequencing data on sorted cell populations; Bottom: aggregated cells within each cluster from sciMET-cap dataset. **c.** Cluster-specific DMRs with Genscript and Genehancer annotations.

### sciMET-cap achieves improved cell type clustering in human cortex with equal read depth

We previously described a high-coverage sciMETv2 workflow that leveraged a linear amplification workflow (sciMETv2.LA) which enables greater library complexity at the expense of increased costs and labor. We had applied sciMETv2.LA to a middle frontal gyrus specimen from a human cortex and produced a total of 927 cells with a median of 1.67 million unique CG sites covered from a median of 5.5 million raw reads per cell for a combined total of 6.98 billion raw sequence reads. This high-depth dataset was combined with two additional preparations using the sciMETv2.SL workflow for a combined total of 2,546 cells which were used to identify 14 clusters that were assigned to cell types, providing high-quality cell type classifications for each cell [15]. Cluster identification was performed using 250 kbp tiling windows and assessing non-CG methylation (CH), a context that is only present in abundance in neurons within brain tissue and is highly neuronal subtype-specific [24]. To enable a baseline assessment of the high-coverage sciMETv2.LA dataset on its own, we re-analyzed the 927-cell dataset using CG methylation levels in target windows as well as CH methylation across 250 kbp windows. This was compared to the full dataset cell type clusters and showed good concordance for the 250 kbp CH methylation analysis strategy with decreased granularity at 10 versus 14 clusters. Evaluation of target window CG methylation permitted a coarse resolution of cell types, identifying only 7 clusters (Fig. S3a). We decided that this high-coverage preparation is an ideal control for comparing sciMET-cap, with ample library to carry out capture on the same input material.

We next performed target capture on the sciMETv2.LA library using standard blockers and washing conditions (cap) and sequenced the library with a total of 339 million raw reads (340M) along with a random selection of the same raw read count from the original non-captured sciMETv2.LA library (noCap) and evaluated performance. Similar to the results of the PBMC experiments, the cap condition produced a median aligned read enrichment of 7.67 with a median of 1.29 unique CG sites covered per raw sequence read versus 0.75 for the noCap condition (Additional file 1,3). We next used the matched read count data to perform windowed methylation assessment, clustering and visualization (Fig. 4a). When leveraging CG methylation across target windows, the capture condition produced 10 distinct clusters that matched well with previously-annotated cell types in contrast to 4 clusters produced by the noCap control (Fig. 4b, S4a).

**Figure 4.**
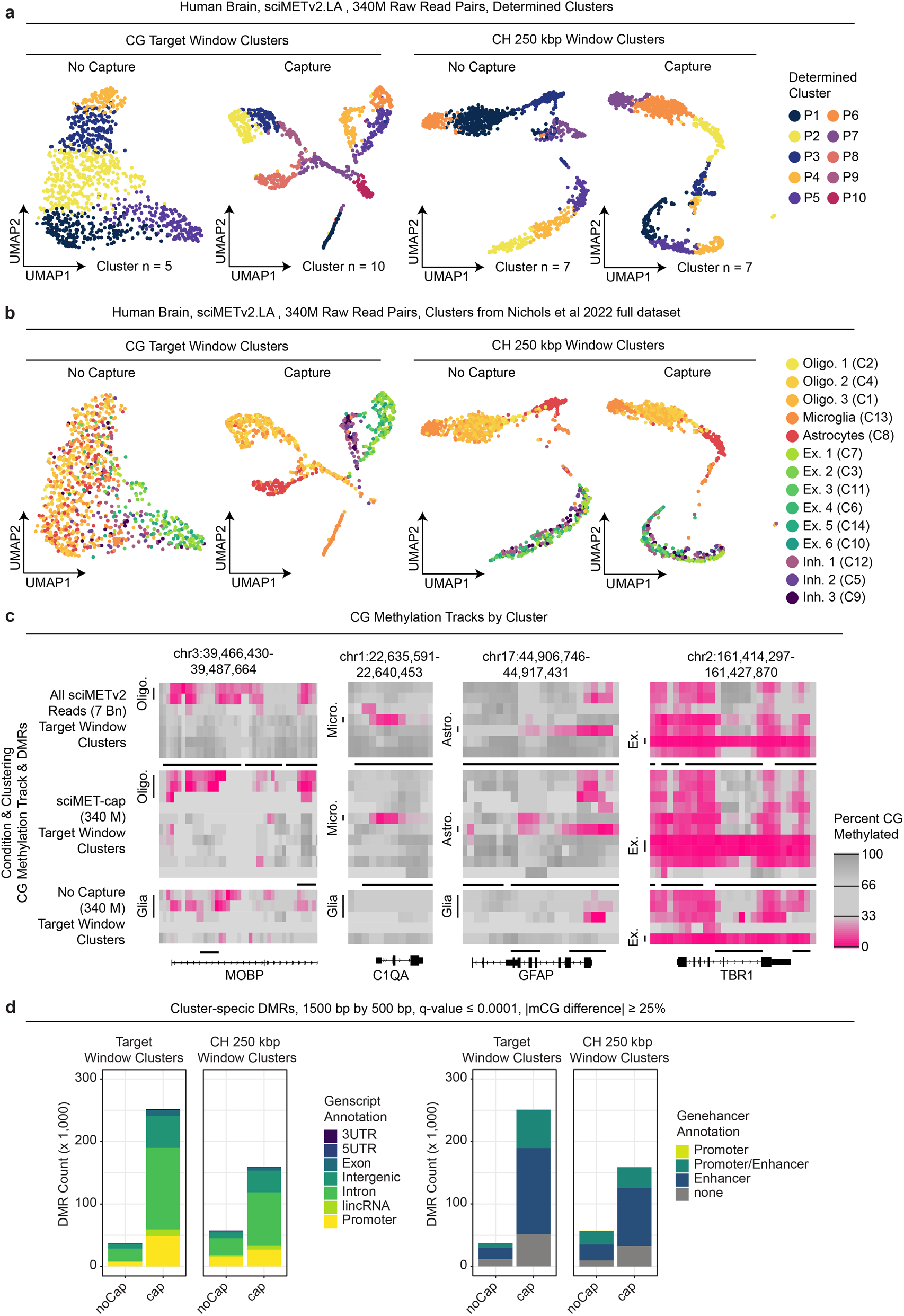
sciMET-cap performance on human brain. **a.** Comparison of sciMETv2 and sciMET-cap using matched read counts, either leveraging CG methylation (left) or CH methylation (right). **b.** UMAPs as in panel a, but colored by previously-defined cell type. **c.** Cluster-specific DMR identification comparison between sciMETv2 and sciMET-cap at matched read depth using either CG or CH methylation for cluster identification. **d.** Genome browser views of CG methylation at marker genes comparing the full sciMETv2 dataset and the downsampled dataset with and without capture.

For the CH windowed analysis, both conditions resolved 7 clusters, suggesting no improvement or detriment in clustering capabilities when incorporating capture. Notably, the ability to discern between excitatory and inhibitory neuron populations was not possible using CH methylation yet CG methylation over target windows produced a clear separation with a single inhibitory and two excitatory clusters (Fig. 4b). In concordance with cell type separation, marker gene methylation levels for the capture dataset were more concordant with the profiles of previously-determined cell types (Fig. 4c). We then assessed the ability to call differentially methylated regions in the downsampled datasets, which produced greater numbers of cell-type-specific DMRs (Fig. 4d), with a greater overlap in called loci with those called from the high-coverage dataset (Fig. S4b).

Finally, we prepared a sciMET-cap library from a separate individual using a single sciMETv2 preparation which was pooled for a single capture and then sequenced on a single P3 flowcell of an Illumina NextSeq 2000 benchtop sequencer; representing a typical experiment with a modest budget. This produced a library containing 6,123 cells and produced 12 distinct clusters using both CG methylation over target windows and CH methylation levels in tiling windows across the genome, with clusters clearly splitting into neuronal and non-neuronal cell types based on global CH methylation levels (Fig. 5a). Marker genes were also examined based on promoter methylation patterns (Fig. 5b), though overall genomic coverage was lower per-cluster due to the reduced read counts present across the large cluster count. This suggests increased sequencing depth may be recommended for tissue types with high heterogeneity, either with higher depth per cell or with more cells at a comparable depth, to ensure high coverage genome-wide. Despite the reduced coverage, 64,774 DMRs were identified that broke down into similar annotations as other datasets (Fig 5c).

**Figure 5.**
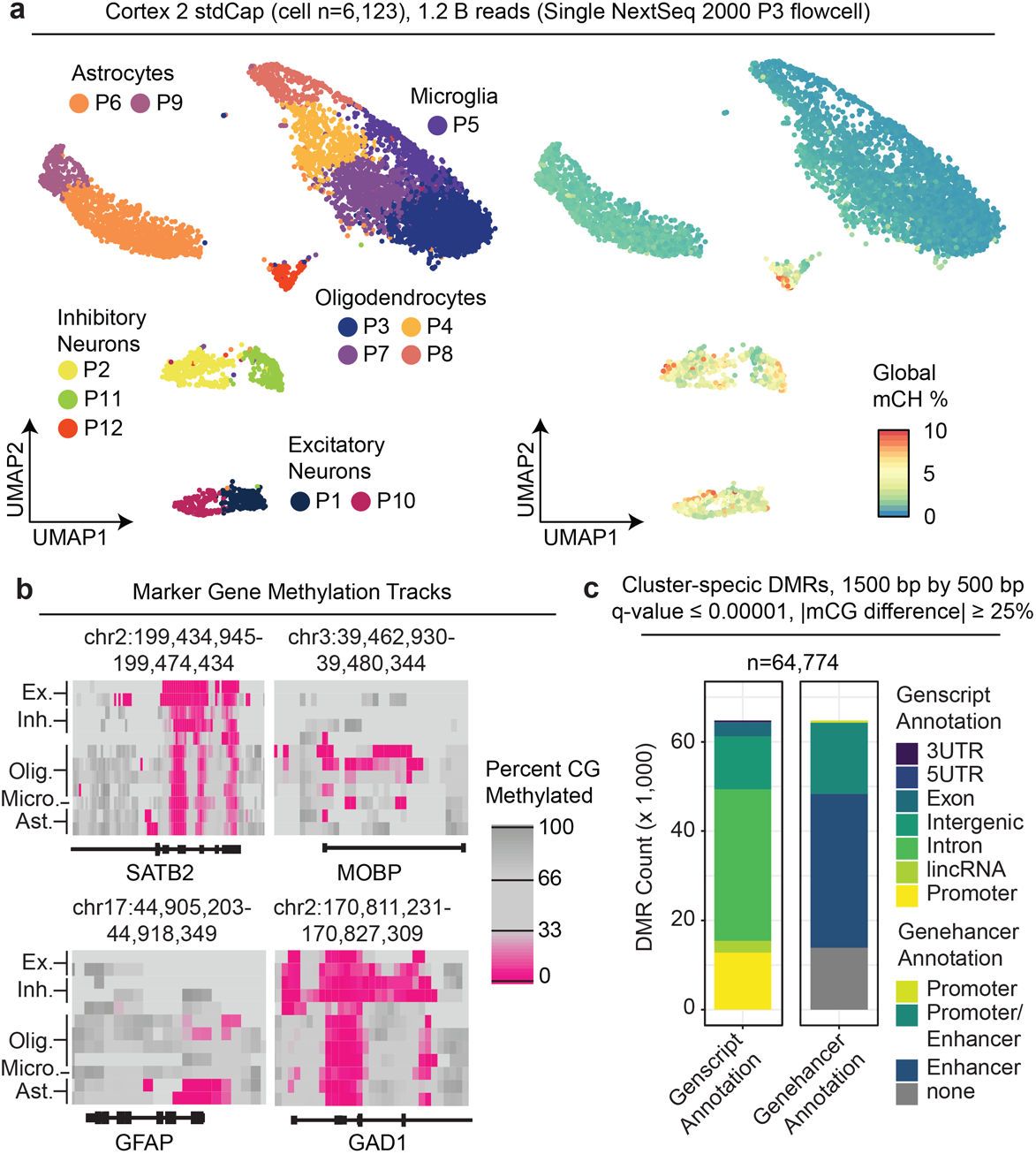
Human brain sciMET-cap using a benchtop sequencer. **a.** UMAP visualization of clusters, annotated by cell type (left) or colored by global CH methylation levels (right) for over 6,000 methylomes produced in a single experiment with a single capture and sequenced on a single flowcell. **b.** Genome browser tracks of CG methylation levels at marker genes for 1,500 bp windows sliding by 500 bp. **c.** Cluster-specific DMRs identified colored by Genscript and Genehancer annotations.

## Discussion

Single-cell DNA methylation analysis has lagged behind that of other molecular properties such as transcriptomic or chromatin accessibility assays. This has largely been due to the lack of high-throughput technologies, with the majority of techniques requiring the deposition of single cells into individual wells for multiple stages of processing, driving up costs and labor. Furthermore, these assays typically cover the entire genome, requiring very high read counts per cell to achieve enough information to perform cell type assignment and clustering. Our recent work detailing sciMETv2 solves the first of these challenges, with the ability to produce thousands of single-cell methylomes relatively inexpensively in a single day by one individual; however, the second challenge of sequence reads requirements remained.

Here, we directly address this hurdle by leveraging off-the-shelf hybrid capture reagents designed for bulk-cell bisulfite or enzymatic conversion-based methylation sequencing with our sciMETv2 technology. We explore several capture conditions using the same PBMC input library which revealed that custom blockers to account for the additional adapter sequences introduced by the tagmentation step do not impact performance. The major factor in on-target enrichment was the wash temperature, with an increased wash resulting in a higher on-target fraction, yet at the cost of reduced library complexity; indicating that standard wash temperatures are favorable to achieve the most information content per cell.

Captured libraries resulted in approximately half of all sequence reads falling within the 125 Mbp target capture region, leaving the remaining half distributed throughout the genome. This enables the use of the enriched on-target regions for cell type assignment followed by aggregation of cells within a cluster to produce full-coverage methylomes. We then evaluated the ability of each condition to capture differentially methylated loci, which produced consistent results across capture conditions, further indicating that the standard wash and blocking conditions are favorable.

Finally, we assessed the capability of sciMET-cap to produce a high-throughput single-cell methylation dataset using only a benchtop sequencing instrument. We produced a sciMET-cap library totaling nearly 6,000 human PBMC cells and were able to clearly identify all major cell type populations from only ∼1.7 billion raw sequence reads. Similarly, we produced a sciMET-cap library on human cortex with over 6,000 cells which we sequenced to ∼1.2 billion raw reads (a single benchtop sequencer flowcell), achieving a similar level of cell type granularity as high-coverage non-capture datasets. This equates to between 200-275 thousand raw sequence reads per cell, which is what we recommended as the target for sciMET-cap for a dataset between 5 to 10,000 cells. While sciMET-cap still requires more sequence reads than is recommended for scRNA-seq (50-100,000) or scATAC-seq (75,000-125,000), it is in a similar range when compared to non-capture single-cell methylation libraries where a target of roughly 2 million raw sequence reads per cell is recommended. We further highlight this reduction in required sequencing depth by producing comparable clustering results from human cortex using sciMET-cap sequenced to roughly 350 million reads versus roughly 7 billion for the same library sequenced without capture.

## Conclusion

In summary, we demonstrate sciMET-cap for target capture of high-throughput single-cell DNA methylation libraries to enrich for regions of regulatory control that exhibit greater cell type variability than the rest of the genome. This enrichment allows for cell type assignment of single-cell methylomes using far fewer raw sequence reads when compared to non-enriched libraries, enabling single-cell DNA methylation experiments to be carried out at a cost point approaching that of single-cell RNA or ATAC (Fig. 6).

**Figure 6.**
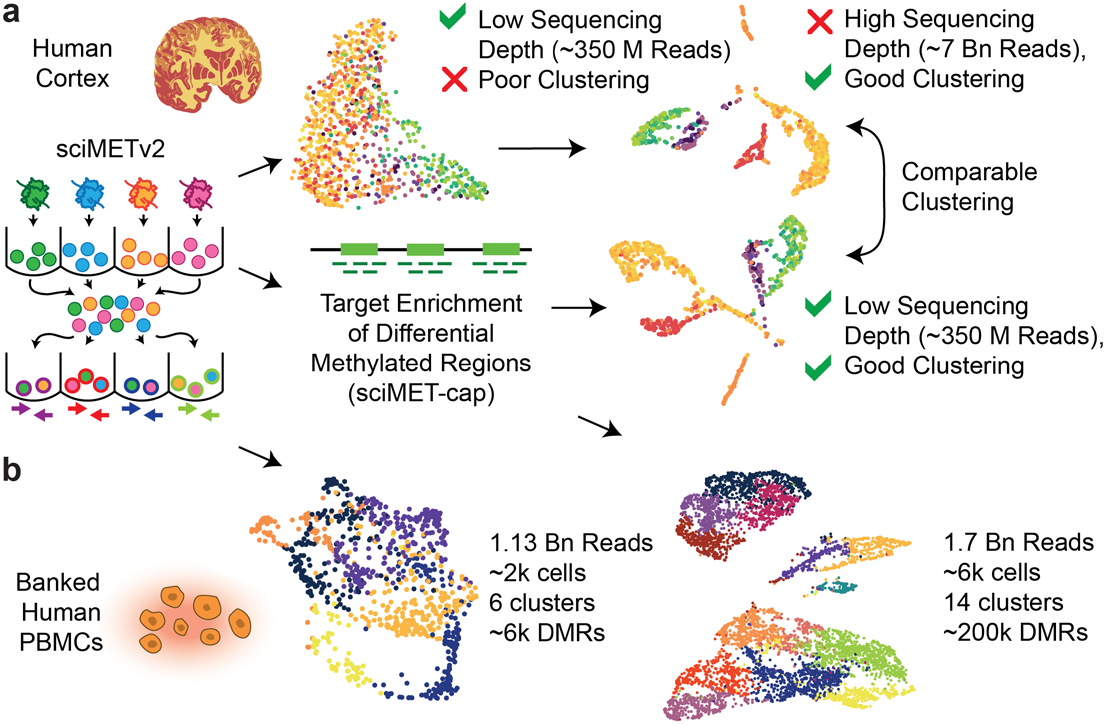
Summary of sciMET-cap performance. **a.** sciMET-cap enables comparable clustering of human cortex single-cell methylation using roughly 350 million sequence reads compared to roughly 7 billion reads without using capture. **b.** Comparison of two experiments on human PBMCs shows the increased power of sciMET-cap to achieve cell type clustering and DMR analysis.

## Methods

### sciMETv2-SL fixation and nucleosome disruption

A detailed protocol for sciMETv2-SL is found in Supplementary Note 1 of Nichols et al. 2022 [15]. Human banked PBMC’s (Peripheral Blood Mononuclear Cells) were obtained from the St. Charles River Labs (PB009C-1/D340161). Nuclei were isolated in NIB buffer on ice for 10’ and spun down at 500xG for 5’ at 4°C. Supernatant was removed and nuclei were resuspended in 200 μL ice-cold NIB. Nuclei were fixed and nucleosome-disrupted according to Nichols et al. 2022 [15] at a concentration of 1 million nuclei per mL. Nucleosome-disrupted nuclei aliquots were then centrifuged for 5’ 500G 4°C and resuspended in 200 μL NIB. Aliquots were combined and nuclei quantified with a hemocytometer This was a safe stopping point and fixed and nucleosome-depleted nuclei could be refrigerated overnight.

### sciMETv2-SL tagmentation

Tagmentation was carried out in a 96-well plate. Prior to nuclei distribution, 4X TAPS-TD buffer was made fresh (1X TAPS = [33 mM TAPS pH = 8.5, Sigma, Cat.T5130], [66 mM KOAc, Sigma, Cat. P1190], [10 mM MgOAc, Sigma, Cat. M5661], [16%DMF, Sigma, Cat. D4551]). Each well contained 5,000 nuclei and 2.5 μL 4X TAPS-TD and enough NIB for a final volume of 10 μL. 2 μL of 5μM Tn5 complexes with methylated C’s, detailed in Nichols et al. 2022 [15] were added. Nuclei were tagmented for 15 min. at 55 °C and placed immediately on ice. All wells were pooled and run through a 40 μm filter. 3 μL of 5mg/ml DAPI was added for flow sorting. 15 nuclei/well were FANS sorted into a plate prepped with 1 μL M-digestion buffer, 0.07 μL reconstituted Qiagen Proteinase K and 0.93 μL H20. Post-sort plates were quick-spun down at 500G, 4°C and then incubated at 50 °C for 20 min.

### sciMETv2-SL bisulfite conversion

Bisulfite conversion was performed as per Nichols et al. 2022 [15], an adaptation of snmC-seq2 [11]. The Zymo Research bisulfite reagents were used for conversion. 1 bottle of CT Conversion Reagent was reconstituted with 7.9 mL M-Solubilization Buffer and 3mL M-Dilution Buffer. After dissolution, 1.6 mL of M-Reaction Buffer was added. 15 μL of the reconstituted CT-Conversion was added to each well of flash-spun post-sort plates. The plates were incubated at 98°C for 8’, then at 64°C for 3.5h with a 4°C hold overnight.

80 μL of M-Binding Buffer was added to each well and then the entire volume of each well was transferred to a 96-well shallow-well Zymo-Spin I-96 plate. The Zymo-Spin plates was attached to a vacuum manifold on the Bravo and flow through was directed into a waste flask and discarded. 100 μL of M-Wash Buffer was added to each well and removed by vacuum flow. The vacuum was turned off. 50 μL of M-Desulphonation Buffer was added to each well. The plate(s) were incubated for 15’ at room temperature. Flow-through was removed by vacuum suction on the manifold and discarded. 200 μL of M-Wash Buffer was added to each well. Flow-through was removed by vacuum suction on the manifold and discarded and plates were spun at 2200 G for 8’ to dry thoroughly. 8 μL Buffer EB preheated to 55 °C was added to each well. This plate was assembled on to a 96-well plate prepared for splint ligation (sciMETv2-SL) containing 1 μL 20ng/ul ET-SSB diluted in SSB dilution buffer. The plates were incubated at 55 °C for 4’. Plates were spun 2200 G for 8’. This is a safe stopping point. Plates can be frozen −20 °C at this stage for at least 3 months.

### sciMETv2-SL splint ligation

The splint ligation plate was heat-shocked at 95 °C for 3’ and iced for 2’. 1 μL of 0.75 uM pre-annealed P5 adapter was added to each well. Ligation master mix was set up at room temperature and 5.2 μL added per well (ligation master mix = 1.63 μL 50% PEG 8000, 0.97 μL 1,3 propanediol (Millipore 807481), 0.75 μL SCR Buffer, 0.6 μL 0.1M DTT, 1 μL 0.1M ATP (ThermoScientific R0441), 0.125 μL T4 PNK (10,000 U/mL, NEB), 0.125 μL T4 DNA Ligase (2,000,000 U/mL, NEB). The addition of 1,3 propanediol was an improvement to the previously published protocol (Kapp et al. 2021, Nichols et al 2022). The plate was shaken at 3,000 rpm for 30s and spun down. It was then incubated at 37 °C for 45’, then 65 °C for 20’ to inactivate the ligase. The plate was spun down again frozen overnight or else indexed as follows.

### sciMETv2-SL indexing PCR

To each well of the splint ligation plate we added 10 μL 5X VeraSeq GC Buffer, 2 μL 10 mM dNTP’s, 1.5 μL VeraSeq Ultra polymerase, 24 μL H20, 0.5 μL 100X EvaGreen, 1 μL 10mm TruSeq i5 and 1 μL 10mM TruSeq i7 primers. The qPCR was programmed to run as follows; 98 °C 30s initial denaturation, 98 °C 10s denaturation, 57 °C 20s annealing, 72 °C 30s extension, 72 °C plate read. The plate was pulled when the majority of wells began to plateau. All wells were partially (10 μL /50 μL) pooled, cleaned and quantified. The partial pools were used to test different capture conditions.

### Hybridize capture probes with sci-METv2 library pool

After pooling 10 μl /50 μl of the indexing plate the mass of the resulting library was 2 μg in 15 μl of water or 5mM Tris. We performed two captures (standard and custom blockers) with ∼1 μg of library material each. In a tube we combined 4 μl methylome panel (Twist Human Methylome Panel, Twist Bioscience, 105520), 8 μl Universal Blockers (also known as standard blockers, Twist Biosciences, 100578), 5 μl Blocker Solution (Twist Biosciences, 100578), 2 μl Methylation Enhancer (Twist Biosciences, 103557) and 1 μg of library in a volume of 7 μl in a 1.5 mL Eppendorf tube. For the second reaction the custom block (Twist Biosciences, RWG180402-L) was added as a 2 μl spike-in. Tubes were dried down on low heat in a speed-vac for 15’ and checked every 15’ for about an hour.

A thermal cycler was programmed as follows: 95°C hold/ 95°C 5’/ 60°C hold (lid 85°C). 20 μl of 65°C Fast Hybridization Mix (Twist Biosciences, 104180) was added to tubes with the dried down panel, library and blockers. The mixture was solubilized for an additional five minutes before transferring to a 0.2 mL PCR tube. 30 μl of Hybridization Enhancer (Twist Biosciences, 104180) was added, the tube was pulse-spun and then transferred to the hot thermal cycler. The reaction was hybridized for 16 hrs. to allow for efficient hybridization of the large (125 Mb) methylome panel.

### Binding and Washing of Hybridized Targets to Streptavidin Beads

Before proceeding, Wash buffers were preheated, and streptavidin beads were equilibrated to room temperature and prepared by washing three times with Fast Binding Buffer (Twist Fast Hybridization and Wash Kit with Amp Mix, 104180; Twist Binding and Purification Beads, 100983). The beads were placed in a 0.5mL tube so that in later steps the bead-bound library could be placed in a thermal cycler with the lid down. A thermal cycler was preheated to 63°C and a heat block at 48°C.

The hybridized library reaction was added directly to the streptavidin beads from the thermal cycler and rocked moderately for 30 min. at room temperature. The library-bead mixture was pulsed spun down, and let stand on a magnet rack for 1 min. The supernatant was removed and discarded. All steps took place in a thermal cycler with the Fast Wash Buffer 1 preheated to the desired temperature in 150μl aliquots in 0.2mL strip tubes. 200 μl of Fast Wash Buffer 1 preheated to 63°C or 67°C was added to the streptavidin bead-bound library and mixed by pipetting. The reaction was incubated in the thermal cycler for 5 min. at 63°C or 67°C. The reaction was placed on a magnetic rack, the beads were pelleted and the supernatant was removed and discarded. The was repeated twice more. After the third wash had been added and incubated, the bead-bound library was transferred to a new 0.5 mL tube to decrease off-target background. The reaction was placed on a magnetic rack, the beads pelleted and supernatant removed and discarded.

200 μl of 48°C Wash Buffer 2 was added to the streptavidin bead-bound library and mixed by pipetting. The reaction was incubated in the heat block for 5 min. at 48°C. The reaction was placed on a magnetic rack, the beads were pelleted and the supernatant was removed and discarded. This was repeated twice more. The reaction was pulsed and spun. All supernatant was removed with a pipette and 45 μl of water was added and the beads resuspended. This mixture was incubated on ice and was referred to as ‘slurry’ in the next

### Post-Capture PCR Amplification, Purification and QC

Before continuing we collected the target-capture ‘slurry’ from the previous step, fresh 80% EtOH, DNA purification beads, Equinox Library Amplification Mix 2X, Amplification primers ILMN (Twist Fast Hybridization and Wash Kit with Amp Mix, 104180; Twist Binding and Purification Beads, 100983), a Qubit dsDNA Assay Kit for mass quantification of library, and an Agilent Tapestation D5000 or Agilent BioAnalyzer to calculate molarity under specified regions.

A thermal cycler (lid 105°C) was programmed with the following steps: initialization step 98°C 45s x1, [denature 98°C 15s, anneal 60°C 30s, extend 72°C 30s, x8 cycles], final extension 72°C 1 min, hold 4°C. The slurry was mixed by pipetting. 22.5 μl of the slurry was transferred to a 0.2 mL PCR tube. The rest of the slurry was stored on ice and if not used, frozen for subsequent use. To the tube were added 2.5 μl of Amplification Primers ILMN, and 25 μl of 2X Equinox Amplification Mix for a total of 50 μl. The slurry was amplified in the thermal cycler for 8 cycles. The reaction was bead-cleaned and taken up in 30 μl. 1 μl was quantified by Qubit and 1ng/ μl was visualized and quantified on an Agilent Tapestation. The library was sequenced with an Illumina NextSeq 2000 using a P2 200 cycle kit.

### sciMET and sciMET-cap data processing

Raw sequence reads were demultiplexed by matching to a whitelist of expected barcodes allowing a hamming distance of 2 for each specified index using the tool unidex (Supplemental Code, https://github.com/adeylab/unidex), two that were added during PCR (index 1 and index 2) and the third (index 3) that was added during tagmentation which makes up the first 8 bp of read 2. Bases 9-29 of read 2 were then removed, as they contain the transposase mosaic end recognition sequence (ME). Either all reads were used, or reads were downsampled randomly to the specified number (350M raw read pairs for PBMC conditions or 339,177,158 raw read pairs for brain, which was the entire capture dataset) before being carried through subsequent steps.

Reads were trimmed for adapter sequences using sciMET_trim.pl, followed by alignment using BSBolt (v1.5.0) using the wrapper script sciMET_align_BSBolt.pl which runs the aligner with reads 1 and 2 swapped due to the opposite configuration of sciMET adapters compared to traditional bisulfite sequencing adapters and then sorts read by name (cell barcode). PCR duplicates were removed for each cell using sciMET_rmdup_pe.pl and then methyl calls extracted using sciMET_BSBolt_extract.pl which outputs chromosome separated cytosine methylation calls split by CG and CH context which are then sorted using sciMET_sortChroms.pl.

Target enrichment was then calculated by filtering both the pre-duplicate removed and post-duplicate removed bam files to contain only reads that overlap the target capture probe regions by at least 1 bp using bedtools intersect (v.2.29.1).

### Dimensionality reduction and clustering

For each condition we generated a matrix of methylation percentage over a set of windows for each cell using sciMET_meth2mtx.pl using CG methylation over 50 kbp windows for PBMCs or CH methylation over 250 kbp windows for brain, as well as CG methylation over target windows for each sample type. Matrixes were then carried through single value decomposition (SVD) using the irlba (v.2.3.5.1) package in R, calculating the first 50 dimensions. For the single capture human cortex experiment the first 25 SVD dimensions from the target window CG matrix and 250 kbp window CH matrix were merged and used for subsequent clustering and visualization. The SVD matrix was then taken through umap for visualization using the umap R package (v.0.2.10.0) and cluster identification using R-Phenograph (v.0.99.1), which leverages Louvain-based clustering methods.

### DMR analysis

We first generated a matrix of CG methylation over 1,500 bp windows sliding by 500 bp using sciMET_meth2mtx.pl followed by aggregating coverage in each window across clusters using each respective cluster call annotation file with the script sciMET_mtx2methylKitAnnot.pl. The resulting files contain 1,500 bp windows in a format that can be read in as a raw methyl object using the MethylKit toolset (v.1.24.0), but instead are pre-tiled to save on file size and read time. These files were also used as input to sciMET_methylKit2bedgraph.pl which generates a bedgraph file for the sliding windows centered on the middle 500 bp region to aid in visualization using genome browser tools. We next used sciMET_methylKit_aggNOTclusters.pl to aggregate all coverage in all clusters other than the one that will be assessed. This was chosen instead of using the set of all other cluster methylation window sets due to the MethylKit unite function which removes all windows where any single sample has less than the minimum coverage threshold, which we set to 10, which would remove a number of windows from the comparison for all clusters if a single cluster exhibits low coverage. We instead used the aggregate coverage of all clusters except the one being assessed and used MethylKit to identify DMR windows. We then filtered to only include windows with a q-value ≤ 0.0001 and an absolute value methylation difference ≥ 25%. Passing windows were then merged to identify unique DMR regions, and were also assessed for GeneHancer and genCode annotations using sciMET_addAnnotsToDMRs.pl. Euler / Venn diagram plots were generated using the sciMET_compareDMRs.pl script which leverages bedtools merge to identify all overlapping DMR loci across each set of conditions and outputs a set that can be directly plotted using the R packaged ‘eurlerr’ (v.7.0.0).

## Acknowledgements

We would like to acknowledge other members of the Adey Lab as well as members of the O’Roak and Saunders labs at OHSU for helpful discussion and feedback. We also would like to acknowledge Scale Biosciences and Twist Biosciences for providing reagents and materials as well as helpful feedback regarding experiments and analysis.

## Author contributions

A.C.A. conceived the study, performed data analysis, wrote the manuscript and supervised all aspects of the study. S.N.A. performed all capture experiments and helped write the manuscript. R.V.N. performed all sciMETv2 preparations with aid from B.L.O. L.R. aided in data analysis. T.P.B. aided in PBMC data interpretation and aided in study design and analysis. All authors reviewed the manuscript.

## Ethics declarations

### Ethics approval and consent to participate

Not applicable.

---

**Figure S1.**
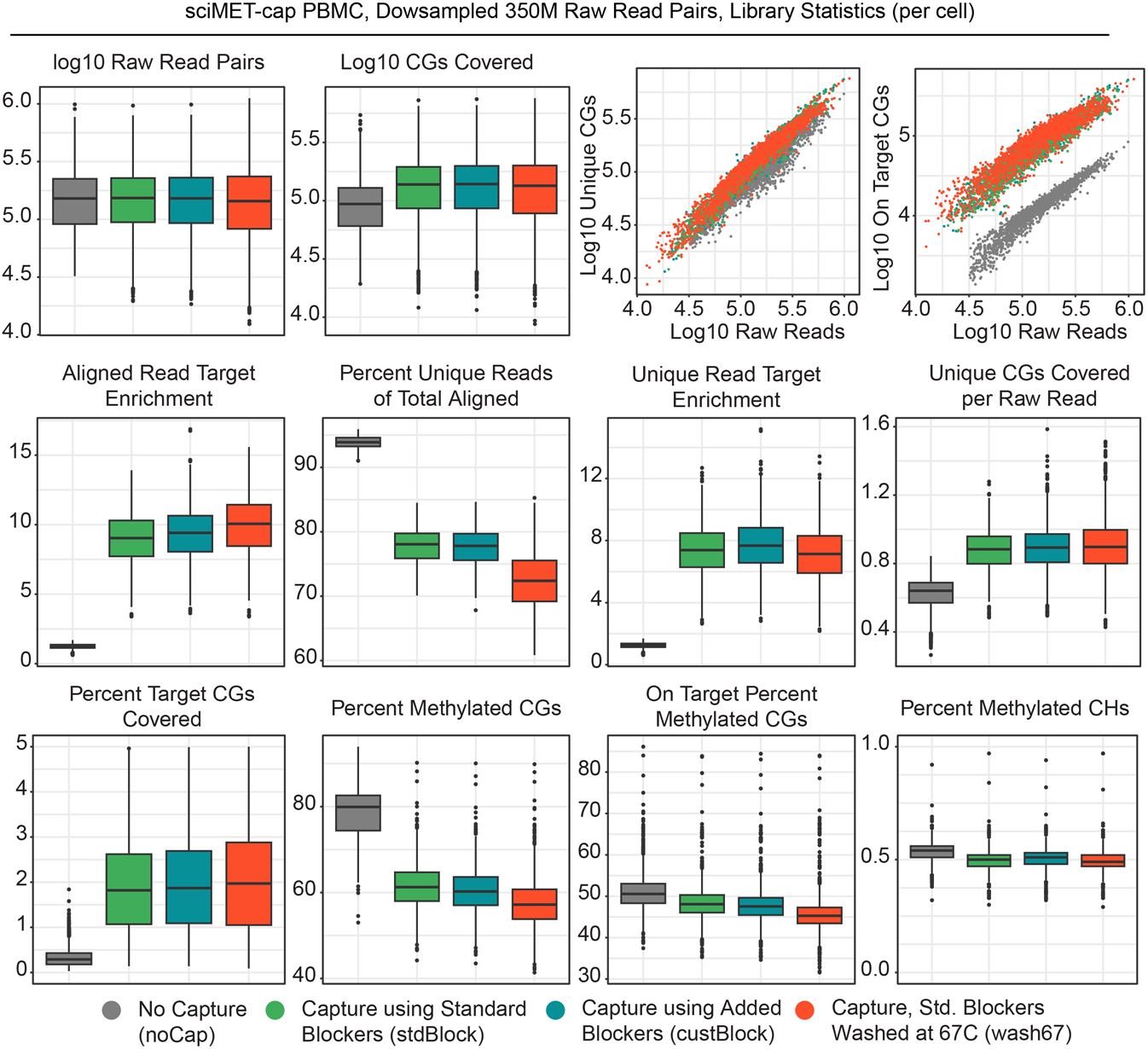
sciMET-cap comparisons. PBMC sciMETv2 and sciMET-cap datasets for four conditions at matched read depth are shown across multiple quality metrics and measurements of sequencing efficiency.

**Figure S2.**
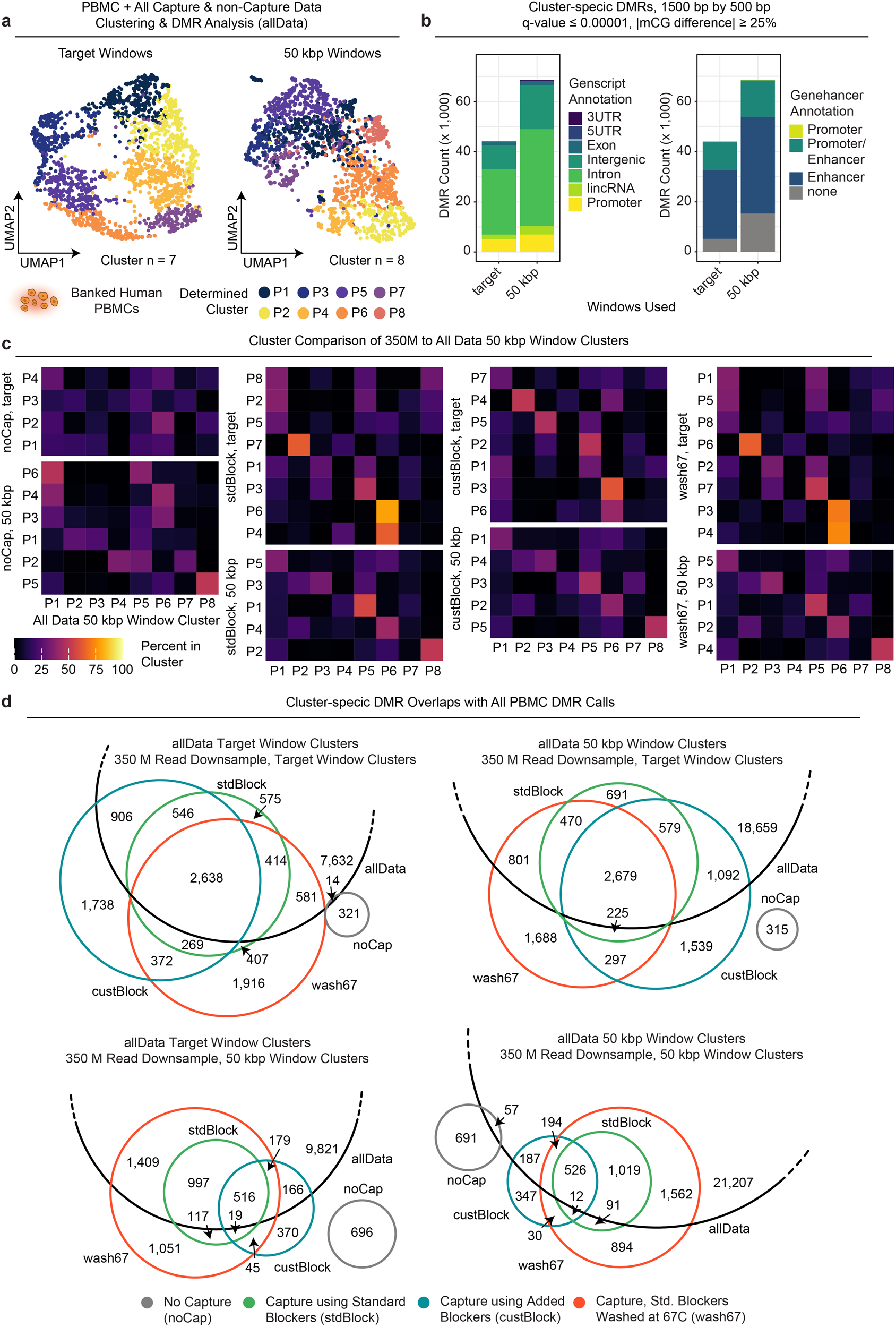
Combined PBMC data and comparisons. **a.** UMAP projections colored by cluster for merged data from all capture and non-capture sequencing runs and conditions using either target windows of 50 kbp tiling windows. **b.** Cluster-specific DMRs identified in the merged dataset with Genscript and Genehancer annotations. **c.** Confusion matrixes of cell assignment across clusters for the 350 million read downsampled conditions compared to the aggregated dataset clusters. **d.** Intersection of cluster-specific DMRs identified across conditions compared with the aggregated dataset DMR calls.

**Figure S3.**
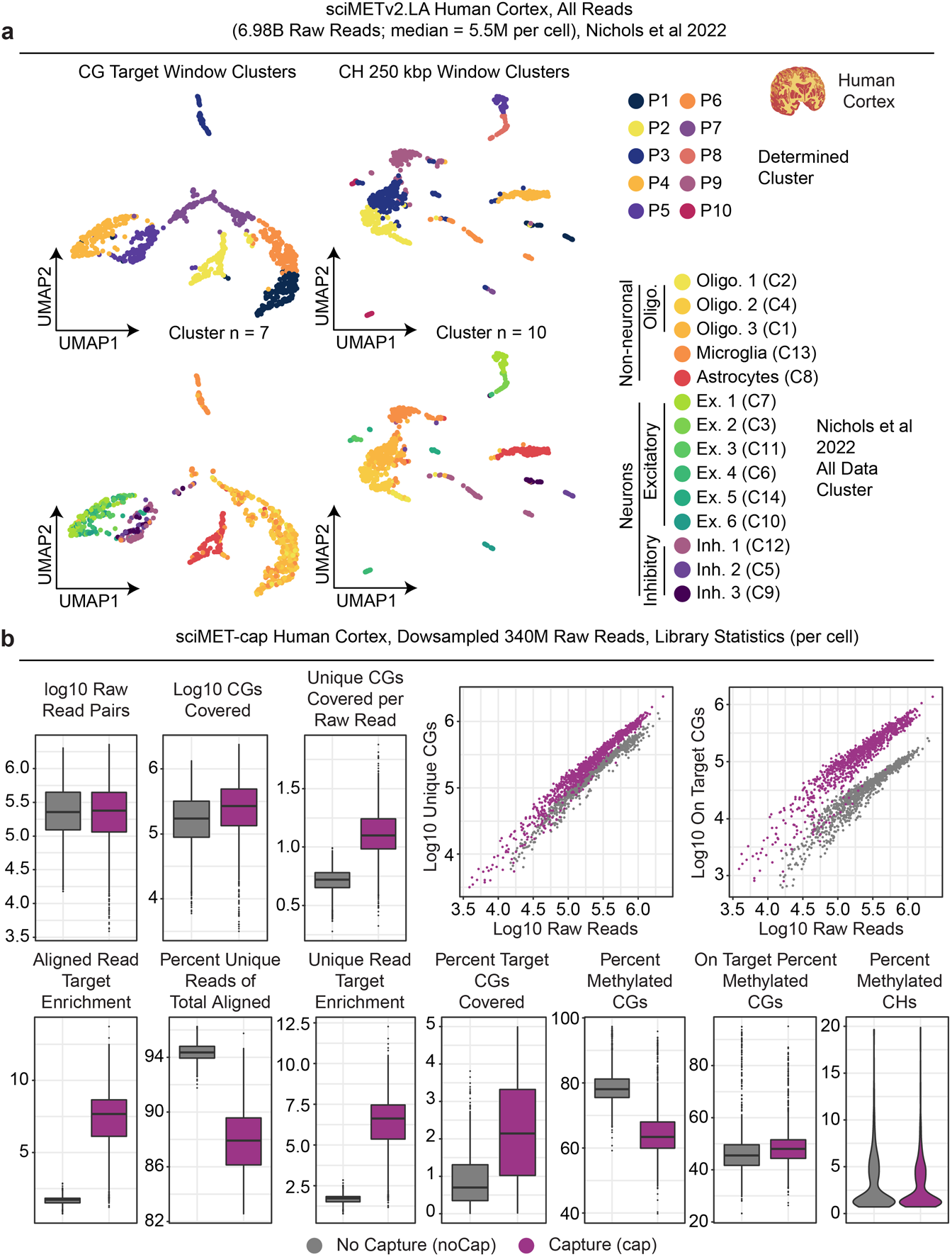
Analysis of sciMETv2 and sciMET-cap on human cortex. **a.** UMAP projections colored by cluster (top) or previously-determined cell type (bottom) for the roughly 7 billion sequence read sciMETv2 dataset. **b.** Comparison of sciMETv2 and sciMET-cap across various quality and sequencing efficiency metrics on matched read count data.

**Figure S4.**
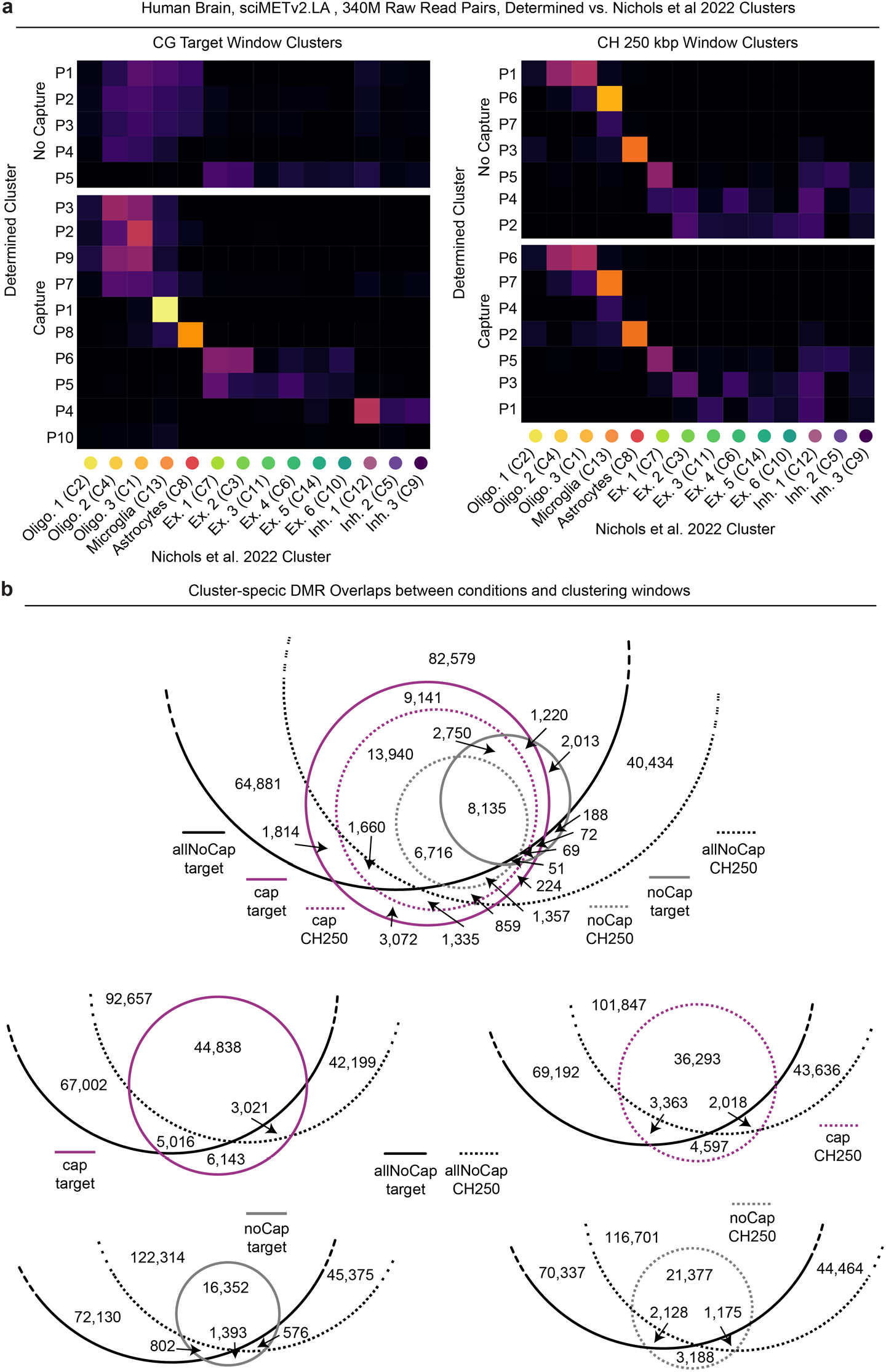
Human brain cluster and DMR comparisons. **a.** Confusion matrixes of called clusters compared to the previously-defined cell types from Nichols et. al. 2022. **b.** Intersection of cluster-specific DMRs identified across conditions compared with the high-coverage non-capture dataset (∼7 billion reads).

